# Temporal fitness fluctuations in experimental *Arabidopsis thaliana* populations

**DOI:** 10.1101/118745

**Authors:** Jinyong Hu, Li Lei, Juliette de Meaux

## Abstract

Understanding the genetics of lifetime fitness is crucial to understand a species’ ecological preferences and ultimately predict its ability to cope with novel environmental conditions. Yet, there is a dearth of information regarding the impact of the ecological variance experienced by natural populations on expressed phenotypic and fitness differences. Here, we follow the natural dynamics of experimental *A. thaliana* populations over 5 successive plantings whose timing was determined by the natural progression of the plant’s life cycle and disentangle the environmental and genetic factors that drive plant ecological performance at a given locality. We show that, at the intermediate latitude where the experiment was conducted, a given genotype can experience different life cycles across successive seasons. Lifetime fitness across these seasons varied strongly, with a fall planting yielding 36-fold higher fitness compared to a spring planting. In addition, the actual life-stage at which plant overwinter oscillated across years, depending on the timing of the end of the summer season. We observed a rare but severe fitness differential after inadequate early flowering in one of the five planting. Substrate variation played a comparatively minor role, but also contributed to modulate the magnitude of fitness differentials between genotypes. Finally, reciprocal introgressions on chromosome 4 demonstrated that the fitness effect of a specific chromosomal region is strongly contingent on micro-geographic and seasonal fluctuations. Our study contributes to emphasize the extent to which the fitness impact of phenotypic traits and the genes that encode them in the genome can fluctuate. Experiments aiming at dissecting the molecular basis of local adaptation must apprehend the complexity introduced by temporal fluctuations because they massively affect the expression of phenotype and fitness differences.

## Introduction

There is overwhelming evidence that local selection pressures have continuously shaped the genetic composition and ecological performance of natural populations [1–3]. But in the context of rapid climate change, greater attention must be devoted to the processes by which species can evolve to adjust their ecological niche [4–6]. The phenotypes that plants deploy in nature, i.e. their timing of germination, their growth rate, their timing of reproduction, are plastically regulated by environmental conditions that vary both temporally and geographically [7]. Much is known about the molecular mechanisms of many plastic plant traits [8–10]. By contrast, we know little about how environmental fluctuations influence the amount of phenotypic variation that is actually expressed in natural conditions, nor do we know how it impacts population levels of fitness variance, an elemental requirement for adaptive evolution [11]. Here we document seasonal variation in expressed plant life cycles and measurements of fitness in *Arabidopsis thaliana*, a plant species that stands as a model for the dissection of molecular modifications underpinning local adaptation [12]. In this species, analyses of genetic variation along altitudinal transects often revealed patterns of local adaptation [13–15]. Patterns of nucleotide variation uncovered the population genetics signatures of local adaptation on a diverse panel of traits [16–20]. Reciprocal transplants of a Swedish and an Italian population suggested phenotypic trade-offs evolved in Scandinavian populations [21,22] and common garden experiments in four different climatic zones highlighted the hundreds of nucleotide variants that associate with local measures of fitness [23]. While these studies reveal the pervasive impact of local selective forces across the range of the species, little is known about the actual ecological challenges populations are exposed to locally and that may lead to this pattern[24]. Because both the environmental conditions and the genetics of specific individuals can vary, the prediction of which plastic traits will be displayed by any one individual and whether it will impact fitness is far from straightforward [25,26]. It is generally understood that local temporal variance of environmental conditions will impact expressed phenotypes and associating fitness levels. Yet, this phenomenon was seldom monitored. At the extremes of the species range, measures of fitness fluctuate across seasons [21,27]. Instead, in climatic regions were summers are sufficiently mild and wet to allow the completion of multiple generation each year, it is unclear how patterns of fluctuations will be affected. Does seasonal variance have cascading effects on expressed life cycles? Does it impact the magnitude of phenotypic and fitness differences manifested by diverse genotypes? Does life history plasticity tend to buffer differences? How does the effect of temporal variance compare with the magnitude of microgeographic or genetic variance?

To answer these questions, we designed an experiment that dissects how environmental factors carve fitness profiles in natural populations at intermediate latitude. We report the monitoring of lifetime fitness in experimental populations over 5 natural successions of plant cycles in conditions differing in soil substrate and water availability. We monitored 4 genotypes, including two genomic backgrounds, one of local and one of distant origin (Col-0 and C24), and the reciprocal chromosome 4 (Chr 4)-introgressions of several polymorphisms previously reported to associate with differential fitness in various locations [23]. Our experiment demonstrates that overall fitness results from complex interactions between seasonal oscillations, soil composition, genomic background and genotypes, leading to marked fluctuations in the phenotypic and fitness differences expressed at a given locality under natural conditions.

## Material and methods

### Genetic plant material

*Arabidopsis thaliana* is an annual weed growing across a broad geographic range [28]. The fruits it produces are called siliques, a character of the Brassicaceae family. In this study, we monitored field performance of two wild type genotypes, Col-0 and C24. Genomic information for these two lines suggest that the Col-0 genotype is typical of Western European populations whereas C24 is more related to Southern European genotypes [29]. We further included in our experiment two reciprocal near-isogenic lines (NILs). The two sets of NILs contained either an introgression of chromosome 4, between 11.8 to 13.8Mb, of C24 accession in Col-0 genomic background or a genomic fragment between 5.1 and 15.3Mb of Col-0 accession in C24 background [30]. Our experiment thus includes genomic background and chromosome-4 introgression as two separate genetic factors. In total, Col-0 and C24 differed for 284 of the 867 SNPs reported to associate with local survival and seed production in *A. thaliana* [23]. Of those, 25 (8%) and 38 (13%) SNPs mapped to the introgressed fragments in Col-0 and C24 backgrounds, respectively. They associated with enhanced survival in Finland and silique number in Spain for the Col-0 genotype [23].

### Field plantings

In this study, we monitored germination, flowering time and fitness of the 2 wild type genotypes, Col-0 and C24 and their reciprocal Chr4 introgressions in the field over five consecutive generations that followed a quasi-natural population establishment dynamics. The field was located in the middle of a flat grass meadow and fenced with iron nets to prevent grazing by rabbits. No vegetation protected it from sun or wind. We filled large flat pots (30cmx40cmx12cm) with two soil types: Arabidopsis turf soil (Einheitserde Typ Minitray, plus 1kg Osmocote Start per m^3^, Einheitserde Werkverband, Sinntal-Jossa) or a soil mixture enriched in sand (a 1:3 mixture of the turf described above and sand). Pots thus dried faster in the sand mixture compared to turf. Pots were randomized and placed on a large plastic cover that prevented germination from the soil below (Suppl. Fig.1). Each replicate population was prepared in a pot by sowing seeds mixed with 5 g silicon dioxide (Sigma-Aldrich, S5631). Pots had been watered to saturation to facilitate seed adhesion to the soil. After sowing, seeds were left exposed to natural conditions in the field and no watering other than natural rain was done. The first planting was started with seeds fully ripened in the greenhouse. All other plantings were started with seeds collected in the field in the previous planting, with the exception of C24 seeds in planting 4, which were taken from planting 2 (the C24 genotype did not survive winter in planting 3). At complete senescence, plants were collected and dried in paper bags and seeds were harvested. Immediately after harvest, 50 field-collected seeds (taken from a pool of 500 seeds collected from 5 plants per pot) of the two NILs and their wild-type parents were placed in 12 replicate pots, for each of the two soil compositions and 4 genotypes. There were two exceptions to this scheme: planting 1 was started with 200 seeds and conducted in only four replicates (32 pots), and planting 5 was performed with only Col-0 and Col^NIL^ due to limitation in personnel (48 pots). The timing of the different plantings is given in Fig. 1.

**Fig. 1:**
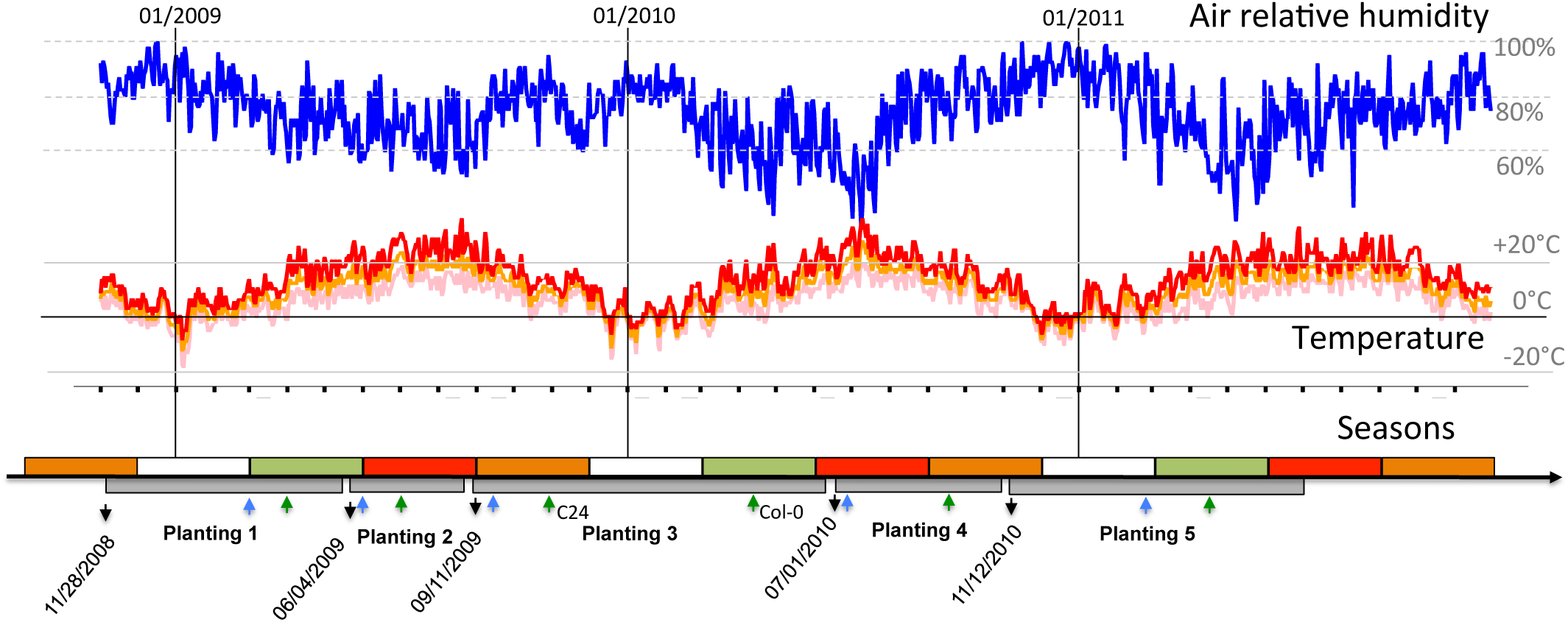
Time plan of the field experiments along five consecutive life-cycles and local meteorological conditions. Meteorological parameters were retrieved from the nearby weather station at airport Norvenich. Relative air humidity is shown in blue, maximum day temperature in red, minimum day temperature in pink and average day temperatue in orange. Black arrows on the lower panels mark the starting date of each planting. Gray boxes mark the duration of each planting. White, green, red and orange rectangles stand for winter, spring, summer and fall, respectively. Blue and green arrows mark the approximate germination and flowering time, respectively. C24 and Col-0 had different flowering times only at planting 3.

For all plantings, seedlings other than *A. thaliana* were immediately removed when they appeared. Our experiment included empty pots to evaluate whether migration occurred between pots. After detecting migrants in control pots, we verified the genotype of each seedling. Individuals with the C24 background display glabrous leaves whereas those of individuals with the Col-0 background are hairy. The genomic background could thus be verified visually. A leaf was collected from each single plant to extract DNA and genotype the individual at Chr4 by PCR with primers mi05 (5’-CTCTTGCTGCGTAGGGTTCCC-3’) and mi06 (5’-ATCTCCCCACTCCCCAATTTT-3’), which generate a length polymorphism at AT4G24415 within the chromosome 4 introgression [31]. We detected 26 heterozygotes in 453 individuals in planting 4, suggesting an outcrossing rate in the field of about 5%. These individuals were removed from further analysis.

### Fitness and phenotype scoring

For the first planting, early, intermediate and late germinants were observed but the exact germination date was not scored. Germination time was scored for plantings 2-5, flowering time (date of the first open flower) was scored for all five plantings. Silique number was counted for a sample of 3-8 randomly chosen plants in each population. In total, fitness was scored for 96, 401, 583, 301 and 691 plants in plantings 1 to 5, respectively (for a total of 2072 plants). In addition, in plantings 2, 3 and 4, the number of seeds was determined for three siliques in each of 3 randomly chosen plants per population. For fruit number and flowering time, an average was computed for each population (e.g. each pot). The average fruit number per pot was taken as an estimate of lifetime fitness. Variation in average seed number per silique did not change the conclusions based on fruit number (not shown). When plants in a pot failed to germinate or survive until reproduction, lifetime fitness was set to zero. To account for variation in germination rates, we included plant density as a covariate in the analyses (see below).

### Statistical analysis

All statistical analyses and figures were performed in R. A generalized linear model (GLM) was used to establish the minimal model explaining phenotypic or fitness variation. For this, we followed the procedure described in Crawley 2005. A first model with average lifetime fruit production as a dependent variable and including, as explanatory variables, the main factors planting, soil type, genomic background and chromosome 4 region, as well as the 2-way, 3-way and 4-way interactions between these factors was run. To take into account variation in fruit number that may result from competition in densely populated pots, the total number of plant in each population was also included as an explanatory covariate. Non-significant interaction terms were removed one by one, from the most complex to the least complex, using the F-test implemented in the R function *drop1.* To control for the non-Gaussian distribution of error, we used a negative binomial distribution of error (implemented in R Package MASS) and tuned its free parameter to adjust the residual deviance to the degrees of freedom of the model. This adjustment was performed at each step of the procedure, until the minimum model was determined. The resulting “minimal” model included the four main factors and the significant interaction terms that remained. Interpreting the effect of a single experimental factor is not straightforward in the presence of interactions between factors, especially when the 4-way interaction is significant. We thus broke down the factors and conducted separate ad-hoc GLM for each genomic backgrounds or for subsets of plantings. The fold-change attributed to experimental factor was calculated by taking the exponential of the log estimates given by the model (GLM with negative binomial distribution of error used the natural log as a link function). This measure corresponds to the odds ratio associated with the factor. The proportion of variance explained by each factor or interaction term, was calculated as the ratio of the deviance computed for a given factor against the total deviance. To compute the fitness fold-change attributed to the chromosome 4 introgression within each combination of experimental factor (planting, soil composition and genomic background), we ran a separate GLM analysis with introgression as a factor nested within a factor called “setting”, which combined soil condition, planting and genomic background in 16 levels. Fitness fold-change attributed to variation in the chromosome 4 region was calculated as described above.

## Results

### Effect of experimental factors on Lifetime fitness

We monitored average fruit (silique) production in experimental *A. thaliana* populations over 5 naturally consecutive plantings by counting the average number of siliques produced per pot (Fig. 1). A generalized linear model with a negative binomial distribution of error adjusted to minimize overdispersion revealed that all factors of the experiment interacted with each other to alter this measure of fitness (Table 1).

**Table 1:** Summary of the deviance table for the minimal generalized linear model testing the effect of planting, soil composition, genomic background and their interaction on lifetime silique productions. Non significant interaction terms were removed sequentially following Crawley 2005. Signif. codes: 0 ’***’ 0.001 ’**’ 0.01 ’*’ 0.05 ’.’ 0.1 ’ ’ 1. Dispersion parameter for Negative Binomial (r=2) family taken to be 0.8980737. Residual deviance: 227.57 for 231 degrees of freedom. GLM final formula: (silinb + 1) ~ Plant Density + Soil Composition + Planting + Genomic background + Chr4 Genotype + Soil Composition:Planting + Soil Composition:Genomic background + Planting:Genomic background + Planting:Chr4 Genotype + Soil Composition:Planting:Genomic background + Planting:Genomic background:Chr4 Genotype + Soil Composition:Planting:Genomic background: Chr4 Genotype.

|  | F (df num, df denom) | Pr(>F) |
| --- | --- | --- |
| Plant Density | 51.3494 ( 1 , 266 ) | 1.03E-11 *** |
| Soil Composition | 21.446 ( 1 , 265 ) | 6.08E-06 *** |
| Genomic background | 47.5303 ( 1 , 264 ) | 5.12E-11 *** |
| Planting | 119.0384 ( 4 , 260 ) | 2.20E-16 *** |
| Chr4 Genotype | 1.7507 ( 1 , 259 ) | 0.1871 |
| Soil Composition:Genomic background | 37.8494 ( 1 , 258 ) | 3.34E-09 *** |
| Soil Composition:Planting | 7.5266 ( 4 , 254 ) | 1.03E-05 *** |
| Genomic background:Planting | 80.7008 ( 3 , 251 ) | 2.20E-16 *** |
| Planting:Chr4 Genotype | 3.1259 ( 4 , 247 ) | 0.01573 * |
| Soil Composition:Genomic background:Planting | 3.6438 ( 3 , 244 ) | 0.01344 * |
| Genomic background:Planting:Chr4 Genotype | 2.8582 ( 4 , 240 ) | 0.02433 * |
| Soil Composition:Genomic background:Planting:Chr4 Genotype | 10.4293 ( 9 , 231 ) | 1.67E-13 *** |

The average number of siliques (fruits) produced in each population was significantly affected by the number of germinants (or survivors) in each replicate population, i.e plant density (p= 1.03e-11, Table 1, Spearman coefficient r = −0.30, p= 3.7e-07, Suppl. Fig. 2). This correlation, however, was present only in experimental plots in which germination and survival rates were high, especially plantings 2, 3 and 5 (Suppl. Fig. 2a). Focusing on planting 2, where both backgrounds could be compared, the decrease of silique production associated with increased plant density was only effective for the Col-0 background (Suppl. Fig. 2b). The C24 background, instead, did not yield high density because of a lower germination rate, especially in sand for the planting initiated in the summer (not shown). In the remaining of the analysis, plant density was included as a cofactor, so that the genetic and environmental effect reported below are independent of population density.

The strongest part of variance was explained by variation across plantings (36% of the variance, p <2.2 e-16, Table 1). Indeed, both highest and lowest values for average silique number per population were observed in planting 3, where most of the experimental populations with C24 genomic background did not survive (Fig. 2A). As a consequence, the second largest effect was controlled by the interaction between genomic background and planting (18%, p<2.2e-16, Table 1). To characterize the details of average fitness variation across plantings for a single genotype reaching reproduction, we focused on the Col-0 background, which completed its cycle in all 5 plantings (Table 2). The log estimate of this analysis revealed that average silique production per plant in planting 3 was 14 times higher than in planting 1, which was itself 2.6 times higher than in planting 5 (Fig. 2B). We thus estimate that, in this experiment, average fruit number per population varied by up to 2.6*14 ~36-fold across plantings for the sole Col-0 genomic background.

**Fig. 2:**
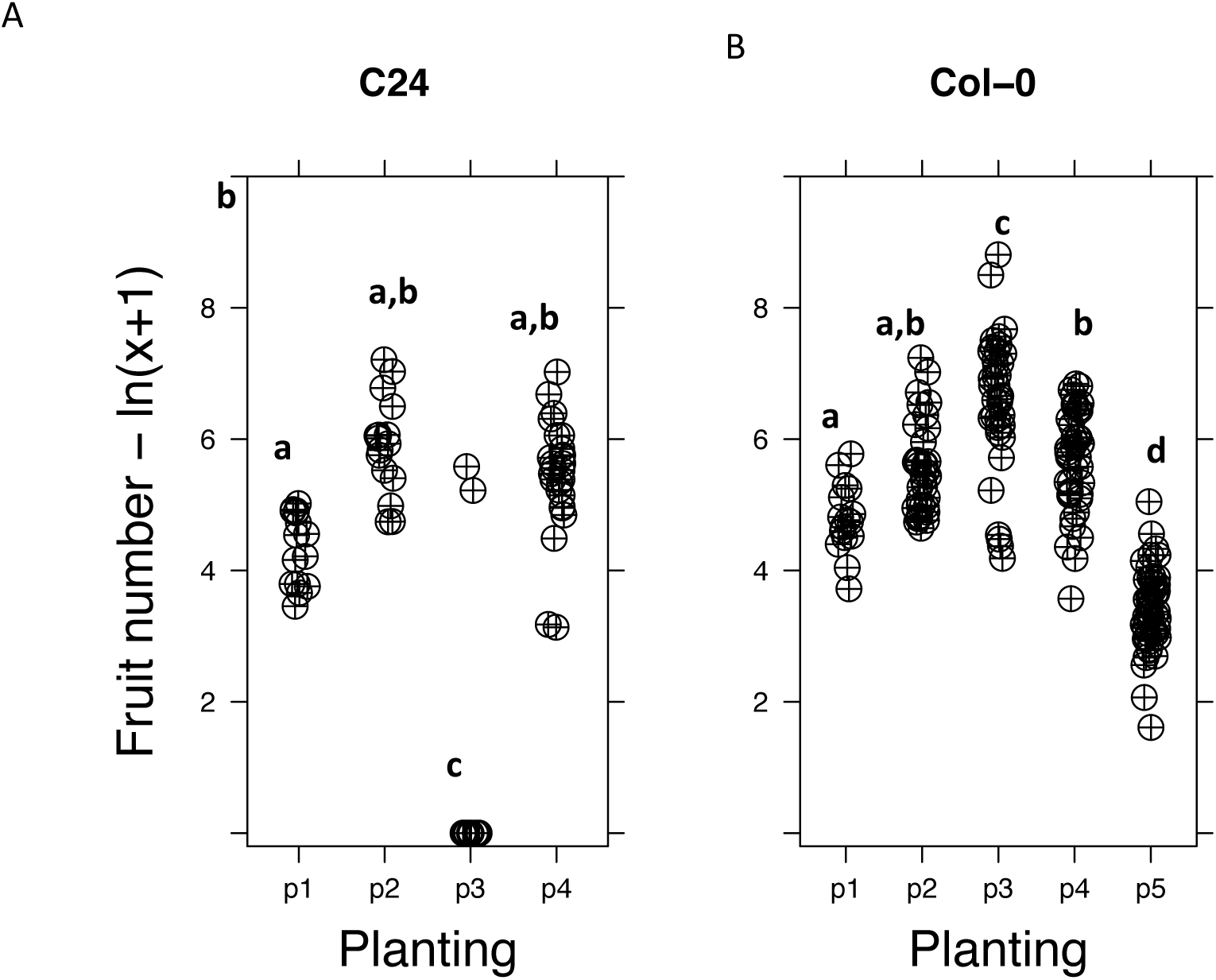
Distribution of total fruit number as a function of planting, quantified as ln(number of fruits +1). A-For individual with the C24 genomic background only, B-For individual with the Col-0 genomic background only. Letters designate significantly different plantings, at *p*=0.01.

**Table 2:** Summary of the deviance table for the minimal generalized linear model testing the effect of planting, soil composition and their interaction on lifetime silique productions of plants with a Col-0 genomic background. Signif. codes: 0 ’***’ 0.001 ’**’ 0.01 ’*’ 0.05 ’.’ 0.1 ’ ’ 1 – Residual deviance: 172.56 for 170 degrees of freedom. Dispersion parameter for Negative Binomial (r = 2.8) family taken to be 0.94596. GLM final formula: formula = (silique number + 1) ~ Plant density + Soil Composition + Planting + Chr4 Genotype + Soil Composition: Planting + Planting: Chr4 Genotype

|  | F (df num, df denom) | Pr(>F) |
| --- | --- | --- |
| Plant Density | 43.4516 ( 1 , 184 ) | 5.25E-10 *** |
| Soil Composition | 45.9537 ( 1 , 183 ) | 1.91E-10 *** |
| Planting | 169.4389 ( 4 , 179 ) | 2.20E-16 *** |
| Chr4 Genotype | 0.4167 ( 1 , 178 ) | 0.519465 |
| Soil Composition:Planting | 4.7902 ( 4 , 174 ) | 0.001097 ** |
| Planting:Chr4 Genotype | 3.1058 ( 4 , 170 ) | 0.016909 * |

As expected soil quality also had a significant effect on silique production (p =6.07e-06, Table 1), with fitness tending to be lower in sand across all plantings, except in planting 3, where growth on sand was advantageous for the Col-0 background (Fig. 3A). Although a relatively small proportion could be attributed to the main effect of genomic background (3%, p= 5.22e-11, Table 1), this factor accounted for a large part of the significant interaction effects with planting and soil. In the three plantings (1, 2 and 4) where the two backgrounds could be compared for fruit production, Col-0 displayed 2-fold greater fitness than C24 in sand, an effect that was not seen in turf (main effect, log estimate 0.85, p= 1.8e-07, interaction with soil, log estimate −0.89, p=9.8e-06, Fig. 3B, Table 3).

**Fig. 3:**
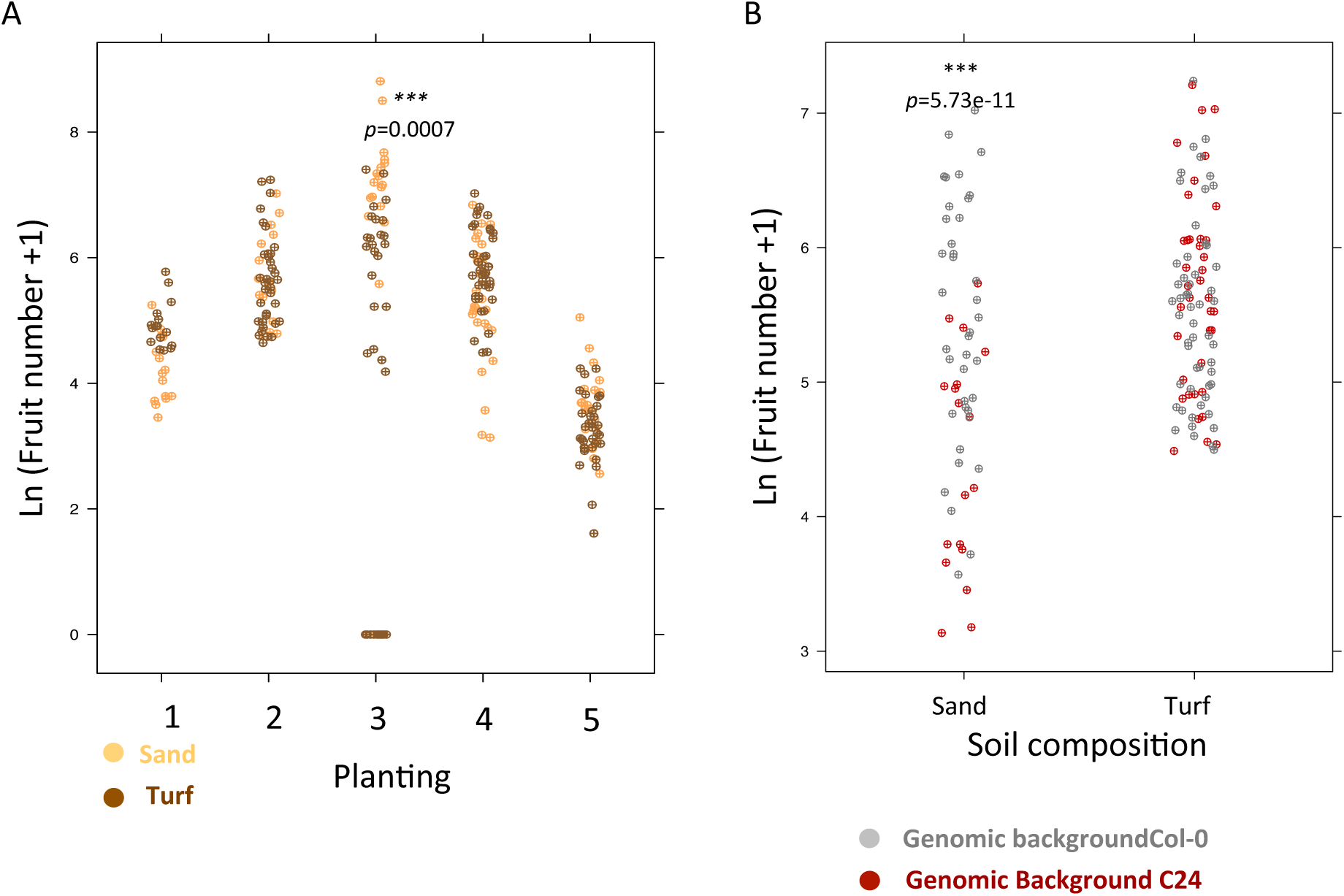
A-The effect of soil on fruit production-transformed as ln(x+1)- changes across generations. In generation 3, plants grown on sand produce significantly more siliques (Fold change=14, p= 0.0007). B-Focusing on planting 1, 2 and 4, where both genomic background survived, reveals that the Col-0 genomic background performs better on sand, whereas growth on turf did not cause strong differences in silique production. ***mark p-values for the subset of conditions where substrate or genomic background had a significant impact on plant performance.

**Table 3:**
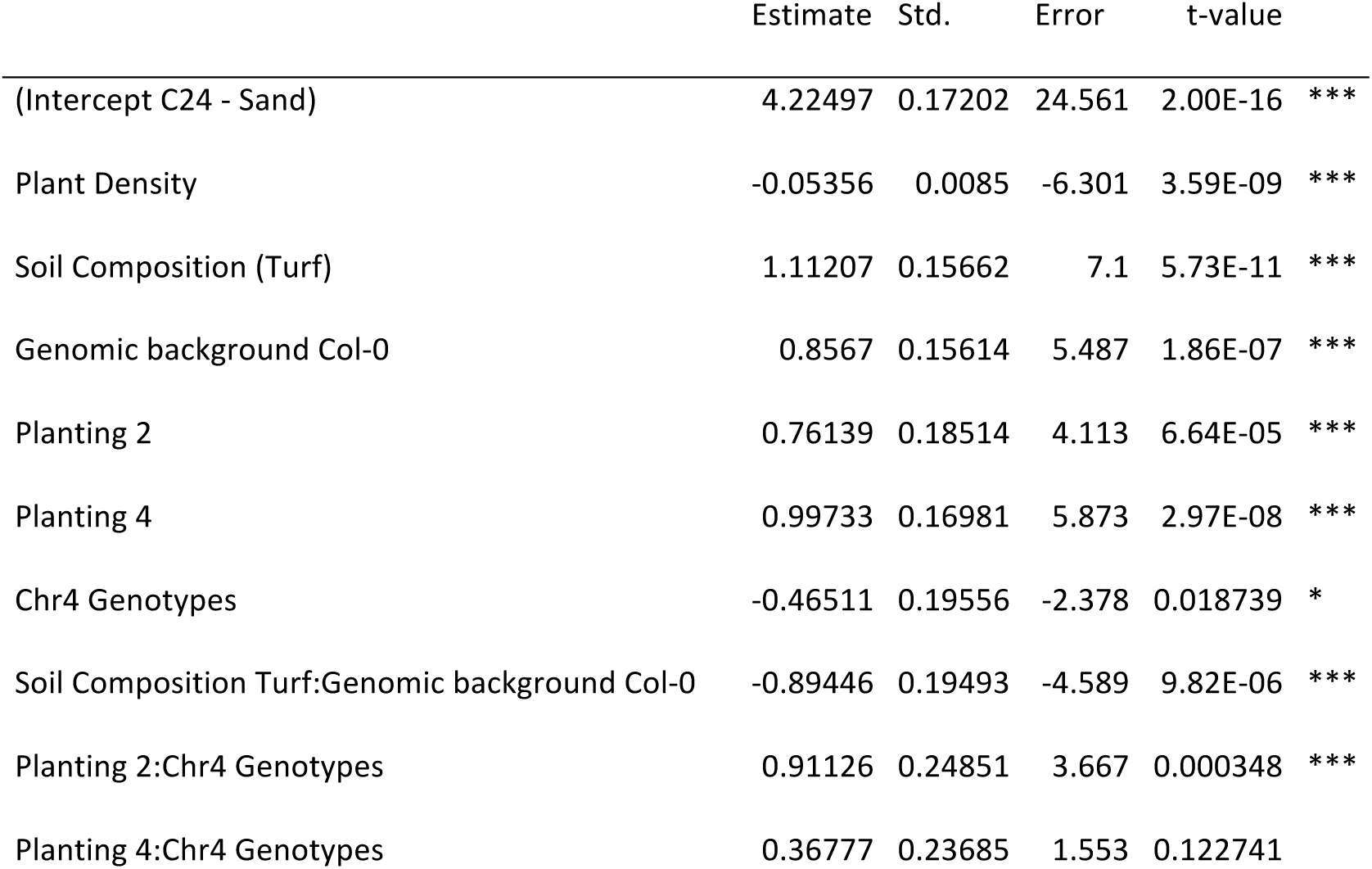
Coefficient estimates for minimal generalized linear model after sequential removal of non significant interaction terms performed on lifetime fruit production (silique number) in planting 1, 2, 4, where both genomic backgrounds survived and reproduced. Signif. codes: 0 ’***’ 0.001 ’**’ 0.01 ’*’ 0.05 ’.’ 0.1 ’ ’ 1 (Dispersion parameter for Negative Binomial (r = 2.9) family taken to be 0.8304712).

The effect of the chr 4 introgression (Tables 1 and 3) was modulated by all factors, including planting, soil and genomic background (F= 10.42, p=1.67e-13, Table 1). The interaction between planting conditions and background also changed significantly the effect of the introgression (F=9.7, p<2.2e-16, Tables 1–2). To investigate which factorial combination revealed the effect of the chr4 introgression, we ran a separate GLM analysis with introgression as a factor nested within a factor called “setting”, which combined soil condition, planting and genomic background in 16 levels (Fig. 4). This analysis revealed that the effect of the introgression was significant for 3 of the 16 settings and its effect was estimated to reverse from a 50% decrease to a 2-fold increase in silique production for the C24 Chr4 allele. Although the introgression of Col-0 into the C24 background was larger than the reciprocal introgression, we detected a significant effect either in both backgrounds or in the Col-0 background only (Fig. 4) suggesting that it is the overlapping portion of the reciprocal introgression that displays fluctuating fitness effects.

**Fig. 4:**
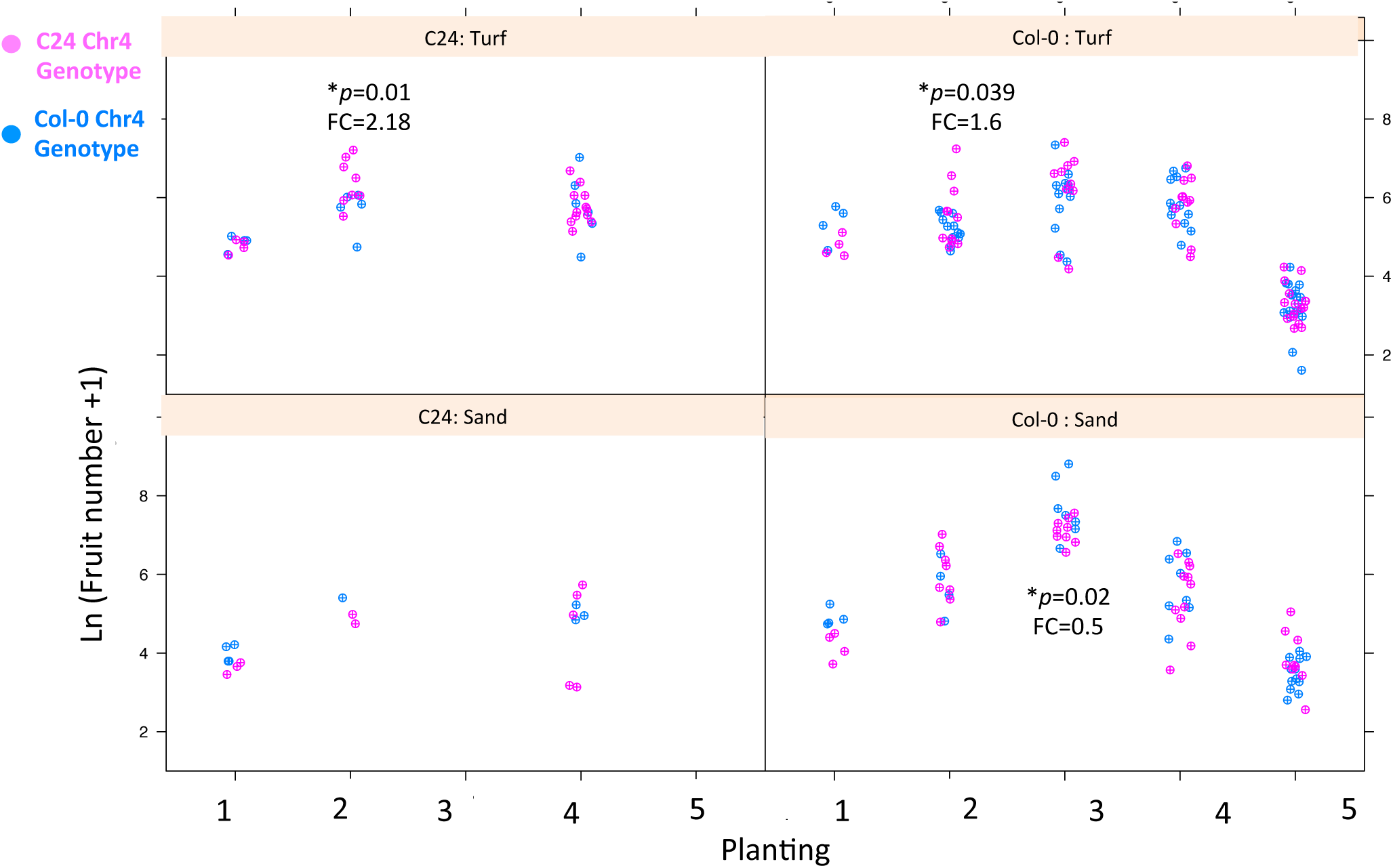
Total fruit number-transformed as ln(x+1)- depends on Chr4 introgression in 3 of 16 combinations of three factors (genome, substrate and planting). *p and FC: p– value and fold-change attributed to the C24 chromosome 4 allele within the factor combination.

The effect of the introgression, when it is significant, is comparable in magnitude to the 2.3- fold increase in silique production per plant of Col-0, compared to the C24 genomic background (Table 3). Thus, in some conditions, the Chr4 introgressed region can reach an effect comparable in magnitude to the average effect caused by the whole genomic background.

### Effect of experimental factors on life-history variation

We observed that plant lifespan varied drastically across the 5 successive plantings (Fig. 1). In the second planting, the life cycle was completed in two months. In planting 3, the life cycle was completed after 8 months and in planting 4, in approximately 4 months. Flowering in planting 4 was terminated in late September (a month later than in planting 2 the previous year), which resulted in a late-fall distribution of seeds for planting 5 and an overwintering at the seed stage.

The duration of vegetative growth is believed to have an important impact on final fitness [27,32]. Yet, in our experiment, we observed no obvious correlation between silique production and the duration of the vegetative life cycle (Fig. 5). Col-0 in planting 3 yielded slightly higher fitness than planting 2 and 4, despite a much longer period of vegetative growth. Yet, in planting 3, a marginally significant correlation between days to flowering and silique number was observed among the Col-0 individuals that survived the winter (Spearman ρ =0.29, p = 0.07).

**Fig. 5:**
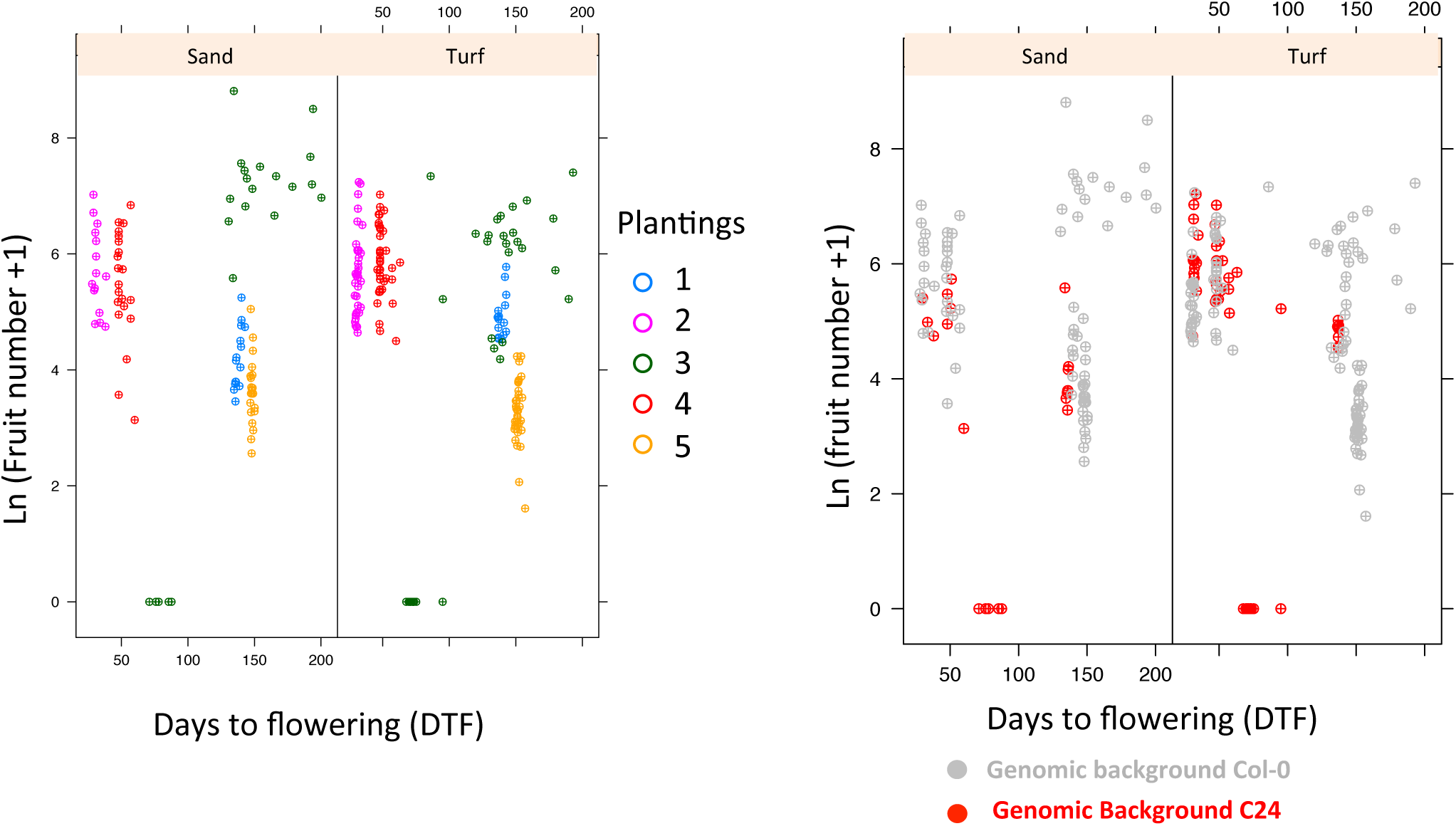
Life time fruit production-transformed as ln(x+1)- as a function of the number of days to flowering in sand and turf. A-Data partitioned by plantings, B-data partitioned by genomic background.

We observed in the 3 overwintering seasons a greater spread in the timing of life-history transition. Planting 1 was started in early December and no germination occurred before spring. Although the precise germination date was not scored for the first planting, we observed that germination was staggered across 1 month. This observation was repeated for planting 5, where germination occurred after 2-3 months (Suppl. Fig.3). For both of these plantings, the date of germination had no impact on the date of flowering: all plants flowered within a week. Our experiment also included another fall planting (planting 3), which displayed germination staggered between 10 and 40 days after seed dispersal. In this planting, the C24 background flowered within a 20-day window (60 to 80 days) in late fall, at a time where the Col-0 background continued vegetative growth. By contrast, the Col-0 background flowered in spring, within a large 80-day window (Suppl. Fig. 3). This spread in the average timing of flowering was observed only in planting 3, the only planting where the two genomic backgrounds showed a significant difference in development (Suppl. Fig. 3).

## Discussion

In this study, we designed a field experiment to quantify the fitness outcome of climate, substrate and genotypic variation on expressed phenotypes and associating fitness levels in a natural setting. Our results reveal that environmental variance experienced over successive growth seasons at a given locality of intermediate latitude is not buffered by phenotypic plasticity. Genotypic differences in phenotype and fitness fluctuate in magnitude across soil conditions and seasons. Seasonal fluctuations experienced at intermediate latitude markedly alter expressed life cycles.

## Seasonal fluctuations in the life cycles expressed by a single genotype

The performance of various *A. thaliana* genotypes in the field has already been reported in several studies [20,21,23,27], but our experiment is the first to monitor the life cycle dynamics naturally deployed by single genotypes. This experiment was conducted along 5 consecutive seasons and included two summer plantings and three fall plantings. From the second season onward, the date of planting was determined by the date of fruit maturation in the field, and the date of germination was determined by the level of dormancy imposed in the field during and after seed maturation. The genotypes were thus allowed to express their natural life cycle. At the temperate latitude of our experimental site, we show that *A. thaliana* completes two cycles per year while the third one is suspended during winter, either at the rosette (planting 3) or at the seed stage (planting 5). Our experiment shows that a given genotype may manifest diverse life cycles in the course of successive seasons, confirming theoretical predictions based on a combination of hydrothermal and photothermal models of germination and flowering decisions[33]. The life stage at which the winter interruption occurred seemed to depend on the timing at which the second generation was finished, either because of variation in the maternally imposed dormancy, the adverse conditions for germination, or possibly the onset of secondary dormancy triggered by low temperatures [34–36].

We further observed that life time fruit production also varied by up to 36-fold across successive seasons. This is likely to be due to the dramatic environmental differences to which genotypes are exposed throughout the successive seasons. Indeed, level of fluctuation for a winter annual cycle replicated over 5 distinct years reported up to 7fold fitness fluctuations [21]. Because overwintering generations tended to reach higher fitness than summer generations, we can further conclude that, at the latitude where this experiment was conducted, variation in characters important for winter survival may be comparatively more crucial for adaptation than those important for completing a life cycle in summer. Nevertheless, since we also observed that the life-history stage of overwintering could change across years, the traits exposed to winter selective pressures also likely fluctuate.

## Seasonal fluctuation in expressed life history differences between genotypes

Based on previous laboratory studies, we expected that the two backgrounds would display markedly different life histories in the field. The genotype C24 was reported to flower 11 days after Col-0 in long day conditions but is a relatively early genotype under short day conditions because of a weak FLC allele [30]. In addition, high ambient temperature (27-30°C) triggers faster flowering in Col-0, but delays flowering in C24, whereas C24 seeds matured under long day conditions at 20°C showed 2 weeks longer primary dormancy than Col-0 (JDM, pers. com.).

Life history decisions such as germination and flowering time are known to be dependent on the environment [27,33,37–39]. Here, we show that phenotypic differences between genotypes can be either masked or exposed across the successive life cycles of a natural setting. Indeed, a major difference in the timing of flowering of Col-0 and C24 was observed only when seeds were sown at the end of August. The late summer time window has been demonstrated to impact the manifestation of flowering time differences controlled by the vernalization pathway [37].

## The Col-0 genotype appears better adapted to the experimental site

The relative fitness of the Col-0 and C24 backgrounds were reported to change markedly across geographical sites [23]. The Col-0 genotype produced 50% more siliques in Halle, Germany, but its fitness was 12% of that of C24 in Norwich, UK, while both genotypes showed comparable fitness in Valencia, Spain [23]. In our experiment, the Col-0 background generally displayed higher fitness than C24 despite temporal seasonal fluctuations. This suggests that it is generally well adapted to the climatic conditions of our field. Its genome is assigned to a Western European *A. thaliana* clade of genotypes originating from natural stands in Germany [29]. The magnitude of the fitness difference nevertheless changed markedly across soil conditions and seasons. A recent study reported that *A. thaliana* genotypes originating from Southern latitudes, which is the case for C24, perform comparatively well at intermediate temperate latitudes [40]. C24, however, did not perform better than Col-0 even in sand where water limitations might have provided it with an advantage. In fact, the Col-0 genotype performed better than C24 in sand, even though plants suffered a 3-week long period of high temperature without precipitation in planting 2 (Fig. 1, Fig. 3). Yet, we also observed that microgeographic heterogeneity in substrate, which is commonly found in natural *A. thaliana* stands, can magnify or mask differences observed between Col-0 and the Southern genomic background C24 (Fig. 3B).

## Rare but radical events counter select inappropriate flowering time

The evolution of gene frequencies in natural populations depends on differential fitness, and therefore on the environmental factors that promote or buffer the expression of this variation[11]. Our experiment shows that although phenotypic differences in the timing of flowering are masked in 3 of 4 seasons, their expression coincided with a severe selection against the early flowering genotype. Flowering however occurred over a broader time window in this planting and a couple of populations (pots) in which a handful C24 individuals which had not flowered, were nevertheless able to regenerate their rosettes and complete their cycle. The transition to flowering might thus have caused the plants’ vulnerability to low temperatures. This experiment therefore depicts relatively rare but radical selective events by which genotypes flowering early in short days might have been counter-selected in regions with sometime severe winter conditions. Such an ecological scenario may explain why local adaptation remains pervasive in this species despite the seasonal fluctuations affecting the expression of adaptive phenotypes that we document [20,23].

## Temporal fluctuations of allelic effects on fitness

Our experiment further shows that seasonal and environmental fluctuations must be taken into account when studying the genomic underpinnings of local adaptation. Extrapolating the evolutionary trajectory of mutations based on their impact on ecological performance in a reductive experimental setting is not directly possible. Whereas the Col-0 background generally performed better over the whole experiment, the Chromosome 4 fragment had effects of changing signs in a small subset of conditions. This small region, which only contains about 10% of previously reported fitness-relevant differences across the geographical range of the species, can have, in some conditions, an effect comparable to that of the genomic background. In fact, interactions between genetic and temporal variation are probably more the rule than the exception. As an example, variation in both plant density and water supply, two environmental factors that are likely to fluctuate over time, has been recently shown to change the effect of a polymorphism impacting water-use efficiency in *A. thaliana* [41].

## Conclusion

The current development of faster and cheaper sequencing methods boosts our ability to investigate the basis of local adaptation in a number of plant species, including rare endemics or aggressive invasive species [42,43]. Annotating how functions encoded in the genome impact ecological performance and fitness seems within reach [44]. Our findings, however, has important bearings for our conception of the specific challenges we are facing for plant (or animal) species that can shift their temporal niches across generations. The likelihood that given genotypes meet the seasonal window where their selective advantage can be exposed will have to be characterized. For this, the genotypic specificities of plastic life history regulation (as e.g. in [45]), the temporal environmental variance, and the scale and impact of micro-geographic substrate heterogeneity will have to be jointly evaluated.

## Acknowledgements

We thank Hildegard Schwitte for her help in quantifying fitness in the 4^th^ and 5^th^ plantings. This research was supported by the Deutsche Forschung Gesellschaft (DFG) in the realm of SFB680, and by the European Research Council with Grant 648617 “AdaptoSCOPE”. The raw phenotypic data presented in this experiment is available upon request.

## Supplemental Figures

**Suppl. Fig. 1:** Photograph of the field experimental set up.

**Suppl. Fig. 2:** A. The effect of plant density in the pot was only strong in planting 2, 3 and 5. For 3 and 5, Co-0 was the only genomic background that survived. B. For planting 2, the C24 background did not perform well, so that density was too low to reveal an impact of competition on total life time fitness.

**Suppl. Fig. 3:** Relationship between days to flowering (DTF) and days to germination (DTG) in each planting for each genomic background. Plantings 2 and 4 had similar germination and flowering time. By contrast, generation 3 showed spread flowering time and generation 5 rather spread germination (spring germination), suggesting that the buffering of germination and flowering differed across planting (and across genotypes). Note that the Chr4 introgression did not impact these differences. In generation 1, the date of germination was not scored, but three cohorts were observed, reflecting a spread of germination over March/April, whereas flowering time occurred within 1-2 weeks.

## References

1. Leimu R, Fischer M. A meta-analysis of local adaptation in plants. Buckling A, editor. PLoS ONE. Public Library of Science; (2008);3: e4010. doi:10.1371/journal.pone.0004010

2. Kawecki TJ, Ebert D. Conceptual issues in local adaptation. Ecol Letters. (2004);7:1225–1241. doi:10.1111/j.1461-0248.2004.00684.x

3. Roux F, Bergelson J. The Genetics Underlying Natural Variation in the Biotic Interactions of Arabidopsis thaliana: The Challenges of Linking Evolutionary Genetics and Community Ecology. Curr Top Dev Biol. Elsevier; (2016);119: 111–156. doi:10.1016/bs.ctdb.2016.03.001

4. Anderson JT. Plant fitness in a rapidly changing world. New Phytol. (2016);210: 81–87. doi:10.1111/nph.13693

5. Chevin L-M, Lande R, Mace GM. Adaptation, Plasticity, and Extinction in a Changing Environment: Towards a Predictive Theory. Kingsolver JG, editor. Plos Biol. (2010);8: e1000357. doi:10.1371/journal.pbio.1000357.g002

6. Hoffmann AA, Sgrò CM. Climate change and evolutionary adaptation. Nature. Nature Publishing Group; (2011);470: 479–485. doi:10.1038/nature09670

7. Sultan SE. Phenotypic plasticity for plant development, function and life history. Trends in Plant Science. (2000);5: 537–542. doi:10.1016/S1360-1385(00)01797-0

8. Jung J-H, Domijan M, Klose C, Biswas S, Ezer D, Gao M, et al. Phytochromes function as thermosensors in Arabidopsis. Science. American Association for the Advancement of Science; (2016);354: aaf6005–889. doi:10.1126/science.aaf6005

9. Legris M, Klose C, Burgie ES, Rojas CCR, Neme M, Hiltbrunner A, et al. Phytochrome B integrates light and temperature signals in Arabidopsis. Science. American Association for the Advancement of Science; (2016);354: 897–900. doi:10.1126/science.aaf5656

10. Andrés F, Coupland G. The genetic basis of flowering responses to seasonal cues. Nat Rev Genet. (2012);13: 627–639. doi:10.1038/nrg3291

11. Wood CW, Brodie ED. Evolutionary response when selection and genetic variation covary across environments. Wiens J, editor. Ecol Letters. 2016. 19:10:1189-1200. doi:10.1111/ele.12662

12. Mitchell-Olds T, Schmitt J. Genetic mechanisms and evolutionary significance of natural variation in Arabidopsis. Nature. (2006);441: 947–952.

13. Luo Y, Widmer A, Karrenberg S. The roles of genetic drift and natural selection in quantitative trait divergence along an altitudinal gradient in Arabidopsis thaliana. Heredity. Nature Publishing Group; (2015);114: 220–228. doi:10.1038/hdy.2014.89

14. Wolfe MD, Tonsor SJ. Adaptation to spring heat and drought in northeastern Spanish Arabidopsis thaliana. New Phytol. (2014);201: 323–334. doi:10.1111/nph.12485

15. Méndez-Vigo B, Picó FX, Ramiro M, Martínez-Zapater JM, Alonso-Blanco C. Altitudinal and climatic adaptation is mediated by flowering traits and FRI, FLC, and PHYC genes in Arabidopsis. Plant Physiology. (2011);157: 1942–1955. doi:10.1104/pp.111.183426

16. Le Corre V, Roux F, Reboud X. DNA polymorphism at the FRIGIDA gene in Arabidopsis thaliana: Extensive nonsynonymous variation is consistent with local selection for flowering time. Molecular Biology and Evolution. (2002);19: 1261–1271.

17. Toomajian C, Hu TT, Aranzana MJ, Lister C, Tang C, Zheng H, et al. A Nonparametric Test Reveals Selection for Rapid Flowering in the Arabidopsis Genome. Plos Biol. (2006);4: e137. doi:10.1371/journal.pbio.0040137.st002

18. Dubin MJ, Zhang P, Meng D, Remigereau M-S, Osborne EJ, Paolo Casale F, et al. DNA methylation in Arabidopsis has a genetic basis and shows evidence of local adaptation. eLife. (2015);4: e05255. doi:10.7554/eLife.05255

19. Brachi B, Villoutreix R, Faure N, Hautekèete N, Piquot Y, Pauwels M, et al. Investigation of the geographical scale of adaptive phenological variation and its underlying genetics in Arabidopsis thaliana. Mol Ecol. (2013);22: 4222–4240. doi:10.1111/mec.12396

20. Hancock AM, Brachi B, Faure N, Horton MW, Jarymowycz LB, Sperone FG, et al. Adaptation to climate across the Arabidopsis thaliana genome. Science. American Association for the Advancement of Science; (2011);334: 83–86. doi:10.1126/science.1209244

21. Ågren J, Schemske DW. Reciprocal transplants demonstrate strong adaptive differentiation of the model organism Arabidopsis thaliana in its native range. New Phytologist. (2012); 194:4: 1112–1122. doi:10.1111/j.1469- 8137.2012.04112.x

22. Ågren J, Oakley CG, Mckay JK, Lovell JT, Schemske DW. Genetic mapping of adaptation reveals fitness tradeoffs in Arabidopsis thaliana. Proceedings of the National Academy of Sciences. (2013);110: 21077–21082. doi:10.1073/pnas.1316773110

23. Fournier-Level A, Korte A, Cooper MD, Nordborg M, Schmitt J, Wilczek AM. A map of local adaptation in Arabidopsis thaliana. Science. American Association for the Advancement of Science; (2011);334: 86–89. doi:10.1126/science.1209271

24. Fournier-Level A, Perry EO, Wang JA, Braun PT, Migneault A, Cooper MD, et al. Predicting the evolutionary dynamics of seasonal adaptation to novel climates in Arabidopsis thaliana. Proceedings of the National Academy of Sciences. National Acad Sciences; (2016);113: E2812–21. doi:10.1073/pnas.1517456113

25. Donohue K, de Casas RR, Burghardt L. Germination, postgermination adaptation, and species ecological ranges. Annual Review of Ecology, Evolution and Systematics 2010. 41: 293–319.

26. Burghardt LT, Metcalf C, Wilczek AM, Schmitt J. Modeling the Influence of Genetic and Environmental Variation on the Expression of Plant Life Cycles across Landscapes. Am Nat. 2015. doi:10.5061/dryad.nv0p1

27. Korves TM, Schmid KJ, Caicedo AL, Mays C, Stinchcombe JR, Purugganan MD, et al. Fitness effects associated with the major flowering time gene FRIGIDA in Arabidopsis thaliana in the field. American Naturalist. (2007);169: E141–E157.

28. 1001 Genomes Consortium. 1,135 Genomes Reveal the Global Pattern of Polymorphism in Arabidopsis thaliana. Cell. 2016. doi:10.1016/j.cell.2016.05.063

29. Nordborg M, Hu TT, Ishino Y, Jhaveri J, Toomajian C, Zheng H, et al. The Pattern of Polymorphism in Arabidopsis thaliana. Plos Biol. (2005);3: e196. doi:10.1371/journal.pbio.0030196.st002

30. Törjék O, Meyer RC, Zehnsdorf M, Teltow M, Strompen G, Witucka-Wall H, et al. Construction and analysis of 2 reciprocal Arabidopsis introgression line populations. J Hered. (2008);99: 396–406. doi:10.1093/jhered/esn014

31. de Meaux J, Hu J-Y, Tartler U, Goebel U. Structurally different alleles of the ath-MIR824 microRNA precursor are maintained at high frequency in Arabidopsis thaliana. Proceedings of the National Academy of Sciences. (2008);105: 8994–8999. doi:10.1073/pnas.0803218105

32. Donohue K, Dorn L, Griffith C, Kim E, Aguilera A, Polisetty CR, et al. Environmental and genetic influences on the germination of Arabidopsis thaliana in the field. Evolution. (2005);59: 740–757.

33. Burghardt LT, Metcalf CJE, Wilczek AM, Schmitt J, Donohue K. Modeling the influence of genetic and environmental variation on the expression of plant life cycles across landscapes. Am Nat. (2015);185: 212–227. doi:10.1086/679439

34. Montesinos A, Tonsor SJ, Alonso-Blanco C, Picó FX. Demographic and genetic patterns of variation among populations of Arabidopsis thaliana from contrasting native environments. PLoS ONE. (2009);4: e7213.

35. Debieu M, Tang C, Stich B, Sikosek T, Effgen S, Josephs E, et al. Co-Variation between Seed Dormancy, Growth Rate and Flowering Time Changes with Latitude in Arabidopsis thaliana. Borevitz JO, editor. PLoS ONE. (2013);8: e61075. doi:10.1371/journal.pone.0061075.s002

36. Postma FM, Ågren J. Maternal environment affects the genetic basis of seed dormancy in Arabidopsis thaliana. Mol Ecol. (2015);24: 785–797. doi:10.1111/mec.13061

37. Wilczek AM, Roe JL, Knapp MC, Cooper MD, Lopez-Gallego C, Martin LJ, et al. Effects of genetic perturbation on seasonal life history plasticity. Science. (2009);323: 930–934. doi:10.1126/science.U65826

38. Chiang GCK, Barua D, Dittmar E, Kramer EM, de Casas RR, Donohue K. Pleiotropy in the wild: the dormancy gene DOG1 exerts cascading control on life cycles. Evolution. (2013);67: 883–893. doi:10.1111/j.1558-5646.2012.01828.x

39. Kronholm I, Picó FX, Alonso-Blanco C, Goudet J, de Meaux J. Genetic basis of adaptation in Arabidopsis thaliana: local adaptation at the seed dormancy QTL DOG1. Evolution. (2012);66: 2287–2302. doi:10.1111/j.1558-5646.2012.01590.x

40. Wilczek AM, Cooper MD, Korves TM, Schmitt J. Lagging adaptation to warming climate in Arabidopsis thaliana. Proceedings of the National Academy of Sciences. National Acad Sciences; (2014);111: 7906–7913. doi:10.1073/pnas.1406314111

41. Campitelli BE, Marais Des DL, Juenger TE. Ecological interactions and the fitness effect of water-use efficiency: Competition and drought alter the impact of natural MPK12 alleles in Arabidopsis. Enquist B, editor. Ecol Letters. (2016);19: 424–434. doi:10.1111/ele.12575

42. Hendrick MF, Finseth FR, Mathiasson ME, Palmer KA, Broder EM, Breigenzer P, et al. The genetics of extreme microgeographic adaptation: an integrated approach identifies a major gene underlying leaf trichome divergence in Yellowstone Mimulus guttatus. Mol Ecol. (2016);25: 5647–5662. doi:10.1111/mec.13753

43. Savolainen O, Lascoux M, Merilä J. Ecological genomics of local adaptation. Nat Rev Genet. Nature Publishing Group; (2013);14: 807–820. doi:10.1038/nrg3522

44. Joly-Lopez Z, Flowers JM, Purugganan MD. Developing maps of fitness consequences for plant genomes. Curr Opin Plant Biol. (2016);30: 101–107. doi:10.1016/j.pbi.2016.02.008

45. Burghardt LT, Metcalf C, Wilczek AM, Schmitt J. Modeling the Influence of Genetic and Environmental Variation on the Expression of Plant Life Cycles across Landscapes. Am Nat. 2015. doi:10.5061/dryad.nv0p1

